# A parasite’s paradise: Biotrophic species prevail oomycete community composition in tree canopies

**DOI:** 10.1101/2021.02.17.431613

**Authors:** Robin-Tobias Jauss, Susanne Walden, Anna Maria Fiore-Donno, Stefan Schaffer, Ronny Wolf, Kai Feng, Michael Bonkowski, Martin Schlegel

## Abstract

Oomycetes (Stramenopiles, Protista) are among the most severe plant pathogens, comprising species with a high economic and ecologic impact on forest ecosystems. Their diversity and community structures are well studied in terrestrial habitats, but tree canopies as huge and diverse habitats have been widely neglected. A recent study highlighted distinct oomycete communities in the canopy region compared to forest soils when taking oomycete abundances into account, in contrast to the homogeneity at the incidence level. It remains however unknown if this homogeneity also leads to a functional homogenisation among microhabitats. In this study, we supplemented functional traits to oomycete canopy and ground communities, which were determined over a time period of two years with a metabarcoding approach. Our results showed that even though most oomycetes occurred in all habitats, a strong discrepancy between the strata and correspondingly the distribution of oomycete lifestyles could be observed, which was constant over time. Obligate biotrophic species, exclusively feeding on living host tissue, dominated the canopy region, implying tree canopies to be a hitherto neglected reservoir for parasitic protists. Parasites highly specialised on hosts that were not sampled could be determined in high abundances in the canopy and the surrounding air, challenging the strict host dependencies ruled for some oomycetes. Our findings further contribute to the understanding of oomycete ecosystem functioning in forest ecosystems.

## 1 INTRODUCTION

Some of the most devastating plant pathogens with worldwide economic and ecologic relevance belong to the Oomycota, protists in the Stramenopiles within the SAR superkingdom (Adl et al., 2019). They comprise several distinct lineages, i.a. the Pythiales, Peronosporales and Saprolegniales (Marano et al., 2014) and occupy ecologically important positions as saprotrophs and severe pathogens. The infamous oomycete *Phytophthora infestans* causes one of the most destructive plant diseases, the potato late blight, and initiated the great Irish famine in the late 1840’s with a million deaths and massive emigration (Mizubuti & Fry, 2006). The ecological and economic impact of oomycetes has led to an increased research interest on their community structures (Robideau et al., 2011; Riit et al., 2016; Singer et al., 2016; Jauss et al., 2020b, 2020a; Fiore-Donno & Bonkowski, 2021), and, correspondingly, their pathogenicity and infection strategies (Rizzo & Garbelotto, 2003; Rizzo et al., 2005; Thines & Kamoun, 2010).

Three lifestyles are described for oomycetes: **Saprotrophic** species are free-living and feed on dead and decaying matter (Lewis, 1973). They occupy key roles in the trophic upgrading of terrestrial, marine and freshwater habitats (Marano et al., 2016). Although saprotrophy is less common in oomycetes, it is believed to be the ancestral state of oomycete nutrition (F. Martin et al., 2016; Spanu & Panstruga, 2017), while the majority of currently described oomycetes are plant pathogens (Thines & Kamoun, 2010). The pathogenic lifestyles include **hemibiotrophy**, characterised by an initial biotrophic phase later turning into a necrotrophic phase after the death of the host (Fawke et al., 2015; Pandaranayaka et al., 2019), as well as **obligate biotrophy**, which comprises species exclusively feeding on living host tissue (Spanu & Kämper, 2010). Even though obligate biotrophic species usually do not actively kill their host, they still damage the host by chlorosis, inflorescence and killing of seedlings, and thus cause severe economic losses (Parkunan et al., 2013; Krsteska et al., 2014; Kamoun et al., 2015).

Oomycete communities are well studied in terrestrial habitats, however, most studies focus on soil and the rhizosphere (Arcate et al., 2006; Esmaeili Taheri et al., 2017; Sapp et al., 2019; Fiore-Donno & Bonkowski, 2021). Recently, Jauss et al. (2020b) characterised oomycete diversity and community composition in tree canopies, which are huge ecosystems containing heterogeneous microhabitats and a large proportion of undescribed diversity (Nadkarni, 2001). Albeit the same oomycetes were present on the ground and in the canopy, communities inhabiting canopy habitats were significantly distinct from soil and leaf litter communities in their abundances. The authors concluded that oomycete diversity in forest ecosystems is shaped by deterministic microhabitat filtering, while a study by Jauss et al. (2020a) could determine air dispersal and convective transport to be the stochastic supplier and distributor of oomycetes among microhabitats and strata. However, the former study only analysed one time point, while the latter study dealing with air samples could show a strong temporal variability in community composition. Accordingly, seasonal variability has been shown to influence protistan communities, to some extent, in several studies (Nolte et al., 2010; Fiore-Donno et al., 2019; Fournier et al., 2020; Walden et al., 2021). For cercozoan communities, Walden et al. (2021) could show annually reoccurring succession patterns in the phyllosphere. This implied not only spatially, but also seasonally structured cercozoan communities in tree canopies, although this was not reflected on a functional scale. If seasonal variation is also reflected in the functional diversity of oomycetes in forest ecosystems, however, remains elusive.

Accordingly, we supplemented functional traits and investigated the seasonal stability of oomycete community composition in forest floors and tree canopies over a period of two years. Our study tackles two hypotheses: (1) Oomycete communities vary not only in their spatial distribution, but also in their seasonal composition, and (2) the deterministic processes leading to differences in community composition between canopy and ground habitats also shape the functional diversity and functional distribution among microhabitats.

## 2 MATERIAL AND METHODS

### 2.1 Sampling, DNA extraction and sequencing

Microhabitat samples were collected in two seasons over a period of two years, i.e. autumn (October) 2017 and 2018 and spring (May) 2018 and 2019 in cooperation with the Leipzig Canopy Crane (LCC) Facility in a floodplain forest in Leipzig, Germany (51.3657 N, 12.3094 E). Samples were obtained and processed as described in Jauss et al. (2020b). Briefly, seven microbial microhabitat compartments related to tree surface were sampled in the canopy at 20-30m height: Fresh leaves, dead wood, bark, arboreal soil and three cryptogam epiphytes (lichen and two moss genera, *Hypnum* and *Orthotrichum*). In addition, two ground samples (soil and leaf litter) were sampled. All microhabitat samples were taken with four replicates, from three tree species with three replicates each. DNA extraction was performed with the DNeasy PowerSoil kit (QIAGEN, Hilden, Germany) according to the manufacturer’s instruction. This procedure was performed on four sampling dates: October 2017 (Jauss et al., 2020b), May 2018, October 2018 and May 2019 (this study). Oomycete-specific PCRs and sequencing were performed as described in Jauss et al. (2020b) with tagged primers designed by Fiore-Donno & Bonkowski (2021); the used primer tag combinations are provided in Supplementary Table 1.

**Table 1:**
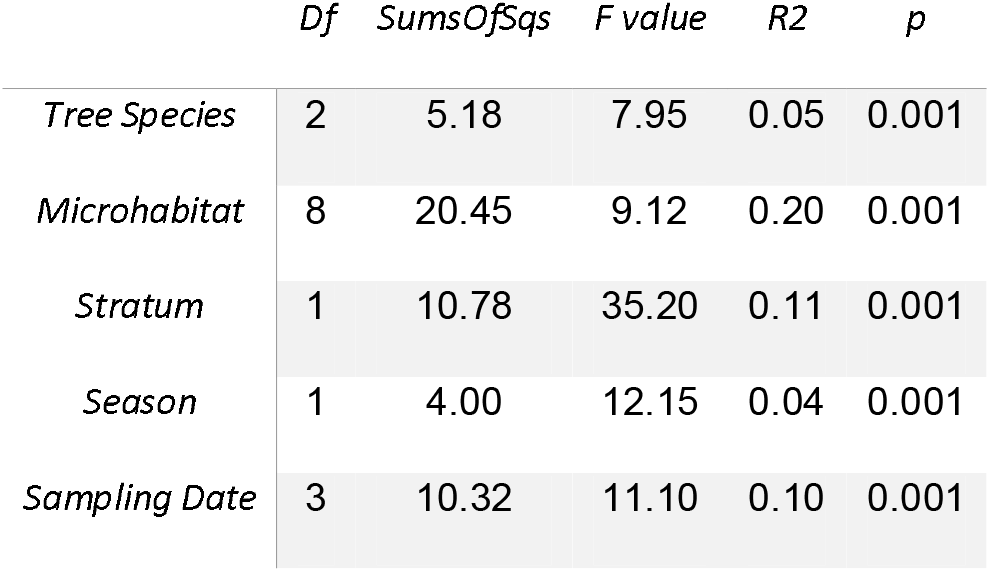
Results of permutational multivariate analysis of variance (permANOVA) from the *adonis* function. Factors were used independently with the default of 999 permutations. *Season* provides the two factors Autumn and Spring, while *Sampling Date* corresponds to the specific time points of sampling, i.e. Autumn 2017, Spring 2018 etc.

|  | *Df* | *SumsOfSqs* | *F value* | *R2* | *p* |
| --- | --- | --- | --- | --- | --- |
| *Tree Species* | 2 | 5.18 | 7.95 | 0.05 | 0.001 |
| *Microhabitat* | 8 | 20.45 | 9.12 | 0.20 | 0.001 |
| *Stratum* | 1 | 10.78 | 35.20 | 0.11 | 0.001 |
| *Season* | 1 | 4.00 | 12.15 | 0.04 | 0.001 |
| *Sampling Date* | 3 | 10.32 | 11.10 | 0.10 | 0.001 |

### 2.2 Sequence processing

Sequence processing and bioinformatics analyses followed the pipeline described in Jauss et al. (2020b). Briefly, raw reads were merged using VSEARCH v2.10.3 (Rognes et al., 2016) and demultiplexed with cutadapt v1.18 (M. Martin, 2011). Primer and tag sequences were trimmed and concatenated sequencing runs were then clustered into operational taxonomic units (OTUs) using Swarm v2.2.2 (Mahé et al., 2015). Chimeras were *de novo* detected using VSEARCH. OTUs were removed from the final OTU table if they were flagged as chimeric, showed a quality value of less than 0.0002, were shorter than 150bp, or were represented by less than 0.005% of all reads (i.e. 368 reads). OTUs were first taxonomically assigned by using BLAST+ v2.9.0 (Camacho et al., 2009) with default parameters against the non-redundant NCBI Nucleotide database (as of June 2019) and removed if the best hit in terms of bitscore was a non-oomycete sequence. Finer taxonomic assignment was performed with VSEARCH on a custom oomycete ITS1 database (Jauss et al., 2020b). The annotation was refined by assigning the species name of the best VSEARCH hit to the corresponding OTU if the pairwise identity was over 95%, OTUs with lower percentages were assigned higher taxonomic levels. Functional annotation was performed on genus level with a custom python script, based on the oomycete functional database published by Fiore-Donno & Bonkowski (2021). Samples with low sequencing depth were removed by loading the final OTU table into QIIME 2 v2018.11 (Bolyen et al., 2019) and retaining at least five samples per microhabitat and 15 samples per tree species per sampling date, i.e. samples with at least 1172 reads. Additionally, the oomycete OTU abundance matrix of air samples from Jauss et al. (2020a) was used for a comparison between tree related microhabitats and the surrounding air from spring 2019, as these samples were taken simultaneously.

### 2.3 Statistical analyses

All statistical analyses were conducted in R v3.5.3 (R Core Team, 2019). Alpha diversity indices were calculated for each sample using the *diversity* function in the vegan package (Oksanen et al., 2019). Non-metric multidimensional scaling was performed on the Bray-Curtis dissimilarity matrix of the log transformed relative abundances (functions *vegdist* and *metaMDS* in the vegan package, respectively), the same matrix was used for a permutational multivariate analysis of variance (permANOVA) with the *adonis* function. Partitioning and visualisation of relative abundances between canopy, soil and leaf litter was performed with the ggtern package (Hamilton & Ferry, 2018). Determination of significantly differentially abundant OTUs was performed with the DESeq2 package (Love et al., 2014). All figures were plotted with the ggplot2 package (Wickham, 2016).

## 3 RESULTS

### 3.1 Taxonomic and functional annotation

We obtained 375 OTUs from 4,262,960 sequences. 77 OTUs (= 20.5% of all OTUs) showed a sequence similarity of less than 70% to any known reference sequence. Plotting the sequence similarity against reference sequences revealed similar patterns as previously described by Jauss et al. (2020b), i.e., many OTUs showed a similarity of 97-100% to known reference sequences, while additional peaks at ~75% and ~85% may indicate hitherto undescribed oomycete lineages (Supplementary Figure 1).

Peronosporales and Pythiales dominated all microhabitats at all sampling events (Supplementary Figure 2). Distribution of functional groups was relatively constant for all four sampling events (Figure 1). Based on OTU presence/absence, the pattern was nearly identical for all microhabitats (Figure 1A-D). Approximately 20% of all OTUs occupied a hemibiotrophic lifestyle, 30% were determined to be obligate biotrophic, only few OTUs belonged to saprotrophic species and the lifestyle of the remaining 50% of OTUs could not be determined, mainly due to low sequence similarities to reference sequences. However, when taking abundances of OTUs into account, the pattern clearly shifted. OTUs assigned to obligate biotrophic species dominated canopy habitats, while ground habitats were more dominated by hemibiotrophic species (Figure 1E-H).

**Figure 1:**
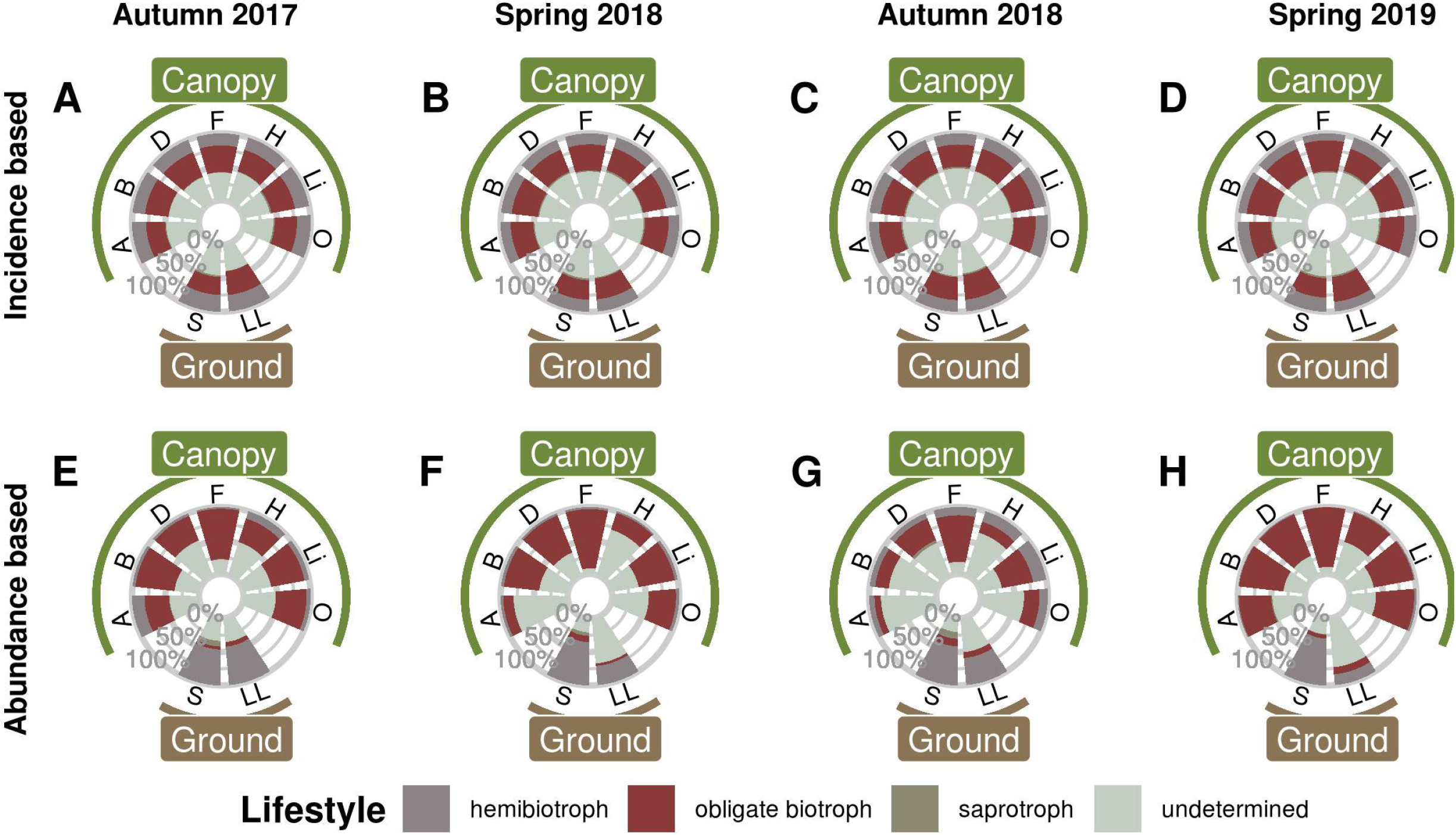
Functional annotation of oomycete OTUs in canopy and ground habitats. (A-D) Distribution of functional groups based on OTU presence/absence, i.e. the proportion of OTUs per Lifestyle. (E-H) Distribution of functional groups when taking abundances into account. A = Arboreal Soil, B = Bark, D = Deadwood, F = Fresh Leaves, H = Hypnum, Li = Lichen, O = Orthotrichum, S = Soil, LL = Leaf Litter

**Figure 2:**
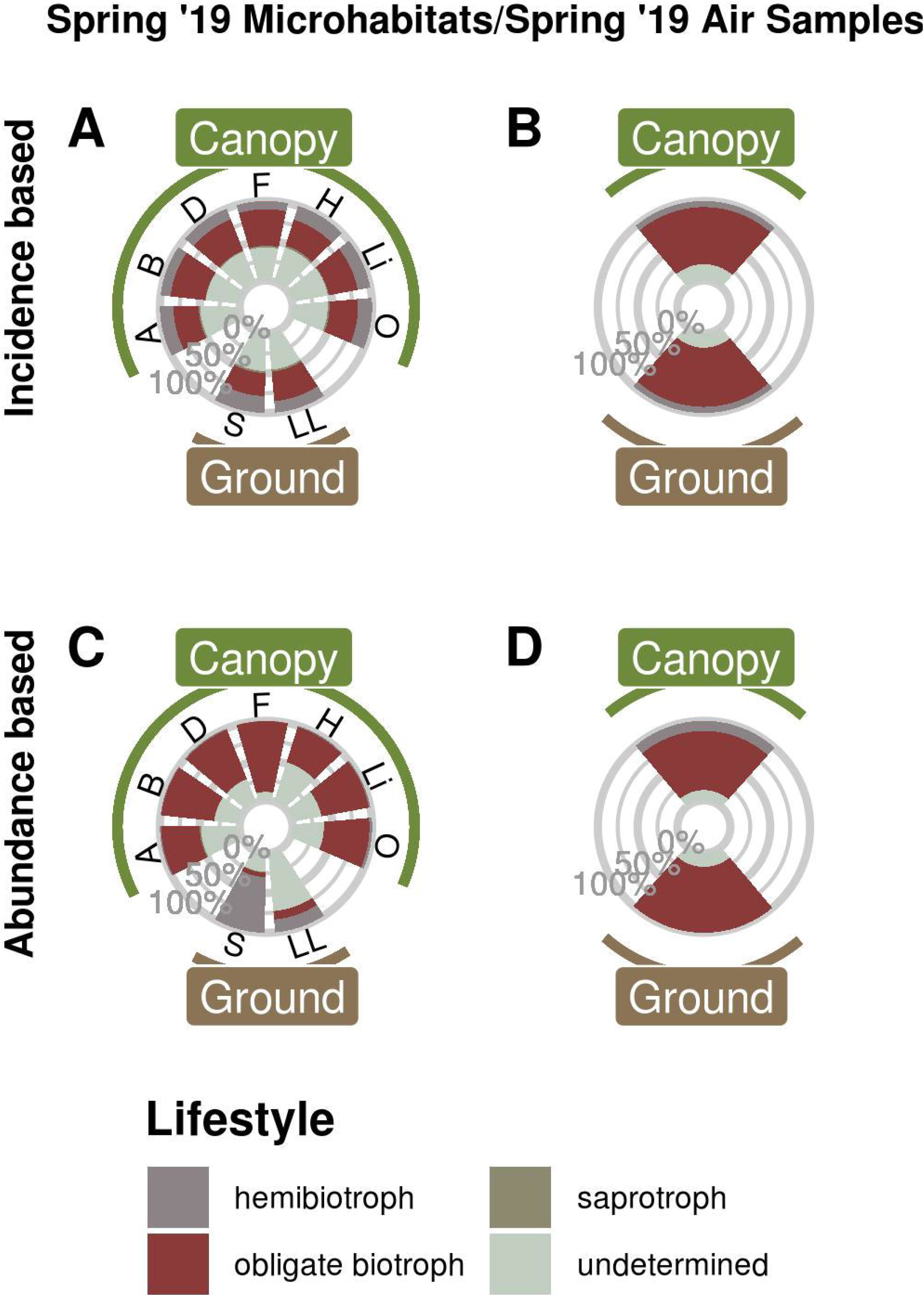
Functional annotation of oomycete OTUs from Spring 2019. Microhabitat samples based on OTU presence/absence (A) and OTU abundances (C) compared to air samples based on OTU presence/absence (B) and OTU abundances (D). For microhabitat abbreviations, see Figure 1.

Comparing the data from Spring 2019 (Figure 1D,H) with air samples previously published by Jauss et al. (2020a) (Figure 2) revealed that the air surrounding canopy and ground habitats was dominated by obligate biotrophic OTUs, irrespective of incidence or abundance.

### 3.2 Abundance partitioning

#### 3.2.1 Partitioning between Canopy, Soil and Leaf Litter

To further determine the distribution of functional groups together with the taxonomic annotation, the relative abundances of each OTU were partitioned for canopy, soil and leaf litter samples (Figure 3). Again, OTUs assigned to obligate biotrophic species dominated canopy samples, while hemibiotrophic species were more evenly distributed or more abundant in leaf litter and soil habitats. Albuginales were almost exclusively present in canopy samples, Peronosporales dominated canopy and leaf litter samples, while Pythiales showed a rather even distribution.

**Figure 3:**
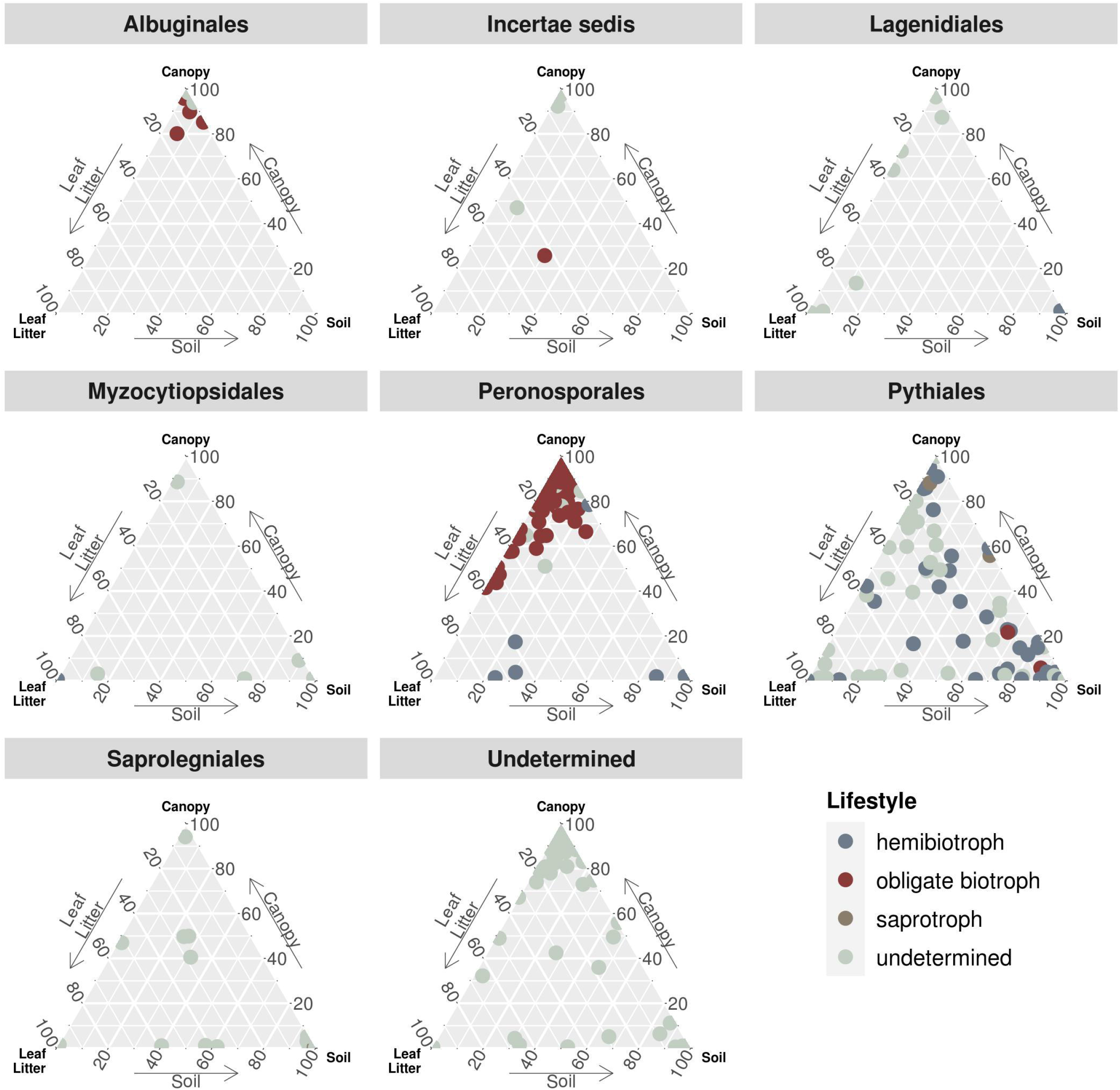
Ternary plot partitioning the relative abundances of OTUs between canopy, soil and leaf litter. Each dot represents one OTU, sorted by taxonomic order and coloured by lifestyle. *Incertae sedis* comprises families and genera not associated with any order, e.g. Lagenaceae or *Paralagenidium*. The order *Undetermined* represents OTUs with sequence similarities of less than 70% to any reference sequence.

The relative abundances of the latter two orders were further partitioned into the four sampling events (Supplementary Figure 3). Abundances of Pythiales were rather homogenous and consistent throughout the seasons, while Peronosporales abundances were more shifted to the canopy region in spring samples. In Autumn 2017, OTUs assigned to the Peronosporales were almost exclusively present in canopy and leaf litter samples, while the distribution in Autumn 2018 was more homogenous.

#### 3.2.2 Differential Abundance Analysis

To determine which OTU abundances were significantly different between the two strata ground and canopy as well as the two sampling seasons spring and autumn, a differential abundance analysis was carried out (Figure 4, Supplementary Figure 4). Within the Peronosporales, this revealed the genera *Peronospora* and *Hyaloperonospora* (obligate biotrophic genera) to be the dominant taxa in canopy samples, while *Phytophthora* (hemibiotrophic) species were significantly differentially abundant in ground samples (Figure 4). For the seasonal effect, more *Peronospora* species were differentially abundant in spring samples compared to autumn samples (Supplementary Figure 4). Within the Pythiales, the genera *Pythium* (hemibiotrophic) and *Globisporangium* (obligate biotrophic) were significantly differentially abundant in ground samples. Most Pythiales, however, could not be determined due to the low sequence similarity to reference sequences.

**Figure 4:**
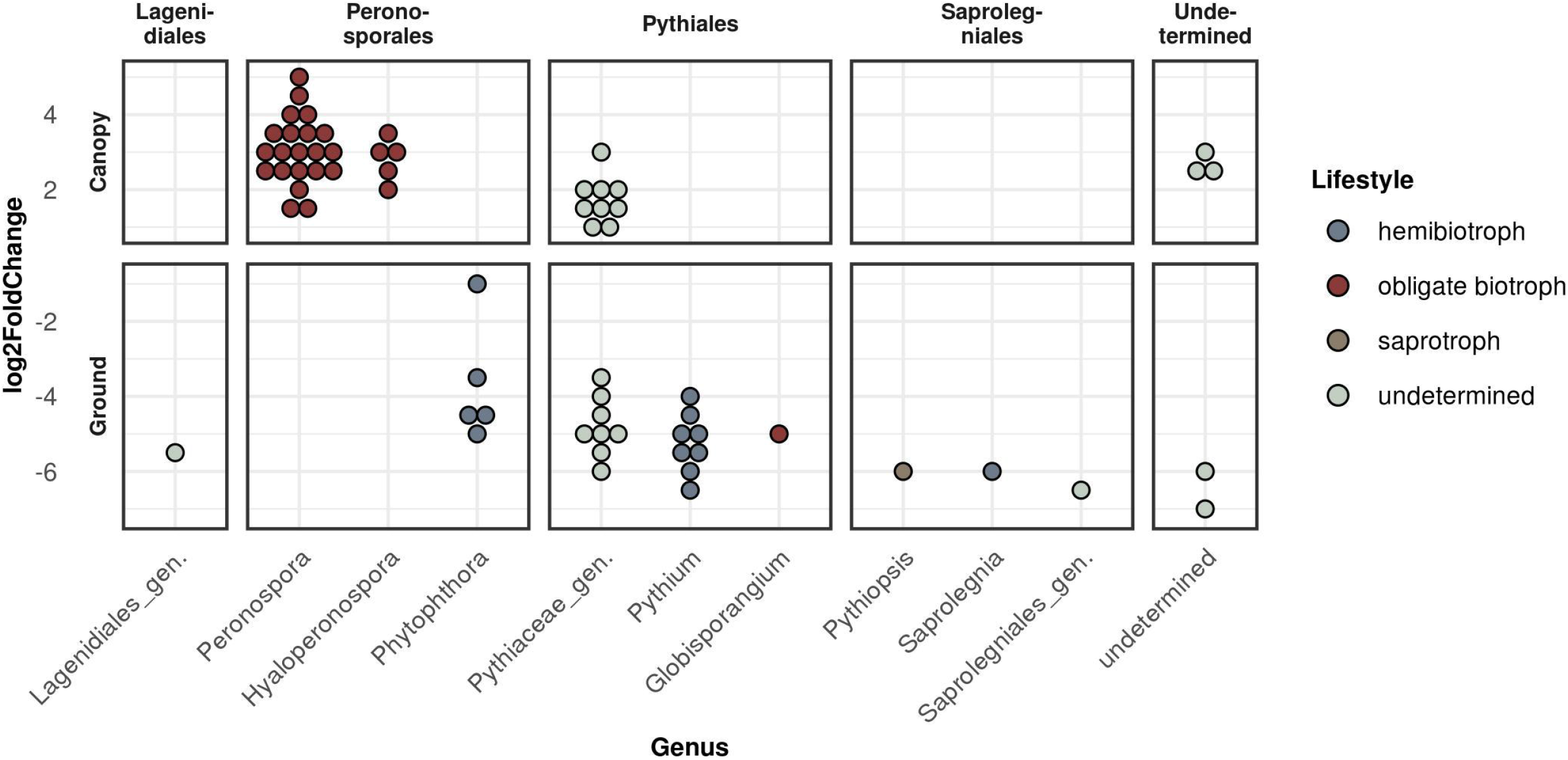
Differential abundance analysis between the two strata canopy (top panels) and ground (bottom panels) sorted by taxonomic order. Each dot represents one significantly differentially abundant OTU grouped by genus. Y-axis (log2FoldChange) gives the measurement of the differential abundance.

### 3.3 Alpha and beta diversity

Despite OTU richness being quite variable among microhabitats, Shannon diversity as well as evenness were high and did not differ between the samplings (Supplementary Figure 5). Beta diversity analyses revealed similar patterns for all seasons as well: the NMDS plot (Figure 5) showed a large overlap of canopy inhabiting communities, which in turn did not overlap with leaf litter and soil communities. This indicated distinct communities inhabiting canopy and ground habitats, respectively, a pattern recurring in all samplings.

Variation in community composition was twice as high among microhabitats (R²=0.20) than between canopy and ground (R²=0.11) or sampling dates (R²=0.10). Tree species (R²=0.05) and season (R²=0.04) explained only a minor fraction of beta diversity (permANOVA, Table 1).

**Figure 5:**
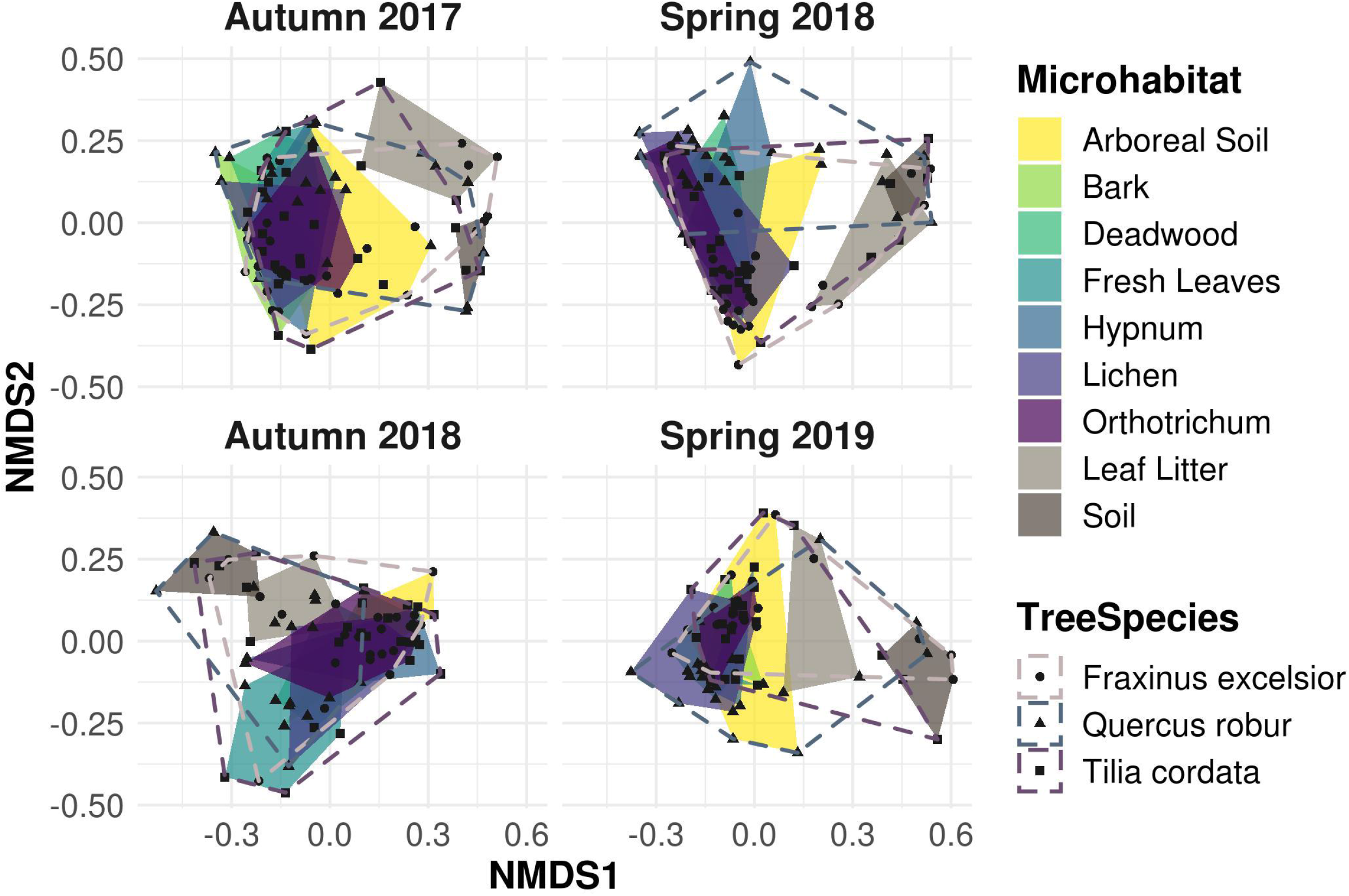
Non-metric multidimensional scaling (NMDS) ordination of Bray-Curtis dissimilarity matrices for canopy and ground microhabitats. Canopy microhabitat communities show a large overlap along all sampling events. Ground habitat communities are strongly separated, indicating unique exclusive communities compared to the canopy region, irrespective of the sampling season.

## 4 DISCUSSION

The most striking pattern of oomycete community composition is the distribution of obligate biotrophic and hemibiotrophic species, with the former dominating canopy habitats and the latter predominantly found in ground habitats (Figure 1). In a previous study, Jauss et al. (2020b) proposed increasing functional diversity instead of increasing species richness with increasing habitat diversity, as most OTUs were shared between all habitats irrespective of specific strata or tree species. Here we supplemented functional traits of the detected OTUs, which revealed that the observed diversity is driven by the lifestyle of the oomycetes. Species occupying a hemibiotrophic lifestyle dominated the two ground habitats soil and leaf litter. Hemibiotrophy is characterised by an initial biotrophic phase, which turns into a necrotrophic phase (Fawke et al., 2015; Pandaranayaka et al., 2019). Oomycetes dwelling the ground habitats are thus capable of feeding on the dead organic matter in the soil, leaf litter and deadwood samples. Deadwood on the forest floor has already been shown to harbour hemibiotrophic oomycetes (Kwaśna et al. 2017a; 2017b). In the canopy, however, deadwood harbours only little hemibiotrophic species, as they are dominated by obligate biotrophic species, like the other canopy habitats. The reason for this might be the high number of obligate biotrophs in the other surrounding canopy habitats as well as in the air (Figure 2). These samples might be overwhelmed by the passive influx of biotrophic species, which are capable of surviving in the other, living, habitats, which would be an interplay between stochastic and deterministic processes for community assembly.

Recent molecular studies analysing oomycete diversity determined similar patterns as reflected in our study, i.e. soil habitats are dominated by hemibiotrophic species, mostly members of the Pythiales (Sapkota & Nicolaisen, 2015; Riit et al., 2016; Fiore-Donno & Bonkowski, 2021). Species of the genus *Pythium* were significantly differentially abundant in our ground habitats. Habitats in the canopy, however, were dominated by the obligate biotrophic genera *Peronospora* and *Hyaloperonospora* (Figure 4). Tree canopies have only recently been subject to studies on microbial diversity (Jauss et al., 2020a, 2020b; Walden et al., 2021; Herrmann et al., 2021), indicating tree canopies to be a hitherto neglected reservoir for parasitic microorganisms. Species of the genus *Hyaloperonospora* are known to be highly host-specific, infecting plant species of Brassicaceae and closely related families (Lee et al., 2017 and references therein). However, none of our sampled trees and microhabitats belong to the Brassicaceae or the order Brassicales. Yet, we observed a high number of reads and OTUs assigned to the genus *Hyaloperonospora* in the microhabitat samples in the canopy as well as in the air samples in both strata, while their number in ground microhabitats is significantly depleted (Figure 4). This indicates a non-random distribution of *Hyaloperonospora* species, as the air as a distribution mechanism should lead to a more or less equal distribution in canopy and ground habitats. Here, they should not be able to survive due to their high host specificity. But the domination in canopy samples implies a capability of survival on hosts they are not specialised on. Thus, we tentatively propose an even less strict host dependency for the genus *Hyaloperonospora* than already suggested (Yerkes & Shaw, 1959; McMeekin, 1960; Dickinson & Greenhalgh, 1977).

The significant differential abundance in the canopy of several undetermined OTUs that can only be assigned to the family Pythiaceae (Figure 4) indicates hitherto undescribed lineages, specialised on the survival in the canopy. Members of the Pythiaceae can occupy all lifestyles, from saprotrophy over hemibiotrophy to obligate biotrophy (Fawke et al., 2015; Marano et al., 2016; Fiore-Donno & Bonkowski, 2021). If the OTUs in the canopy would show an obligate biotrophic lifestyle, it would be in line with observations of the other lineages in the canopy (Figure 1). Yet, the sequence similarity of these OTUs amounts to only ca 80-85% to any reference sequence, thus we only tentatively draw conclusions about their lifestyle.

A common pattern in microbial community ecology studies is a high seasonal variability (Nolte et al., 2010; Fiore-Donno et al., 2019; Fournier et al., 2020; Walden et al., 2021). Oomycete community compositions were in fact slightly, yet significantly distinct for every sampling and correspondingly for every season (Table 1). This pattern is in line with hypotheses proposed by Jauss et al. (2020a), that seasonal variation in air samples drives the community composition in forest ecosystems. The environment, however, then selects the species most adapted to the microhabitat, leading to overall similar community patterns and microhabitat differences for every season (Figure 5). The seasonal changes in microhabitat properties (e.g. temperature, moisture or habitat structure) thus affect all habitats and communities equally. The season itself explained less variance in community composition than the sampling dates (i.e., Autumn 2017 vs. Autumn 2018 etc.; Table 1), suggesting that annual changes do not lead to similar community structures within microhabitats in each season as an annual cycle *per se*, but rather indicate a high temporal variability while preserving spatial diversity. Fournier et al. (2020) observed similar patterns, concluding deterministic niche-based processes in microbial forest soil community assembly. Implications are that ecosystem functioning of oomycete communities is not mainly affected by seasonal fluctuations, but rather by microhabitat identity and, correspondingly, responses of lifestyle to microhabitat filtering (Fiore-Donno & Bonkowski, 2021).

## Conclusions

Both our hypotheses were confirmed in this study: Oomycetes show not only a spatial, but, to a lesser extent, also a temporal variation in their communities. Within the temporal variation however, the spatial variation is preserved, leading to overall similar community patterns for every sampling date. Further, these deterministic processes also shape their functional diversity in forest ecosystems. Our results indicate that tree canopies not only offer numerous distinct habitats to microorganisms, but also serve as a reservoir for parasitic species. Spatial diversity and correspondingly functional diversity drive the oomycete community to a greater extent than temporal diversity. Thus, our findings contribute to future studies on oomycete ecosystem functioning.

## Acknowledgements

The authors would like to thank Rolf Engelmann for his assistance with the field work by operating the canopy crane, as well as the Leipzig Canopy Crane Platform of the German Centre for Integrative Biodiversity Research (iDiv) for providing the site access and allowing us to sample the trees from their field trial.

## Data Accessibility

Raw sequence data have been submitted to the European Nucleotide Archive (ENA) database under the Bioproject number PRJEB37525, with accession numbers ERS4399744, ERS5649966 and ERS5649967.

All figures, codes and detailed bioinformatic/statistical methods used in this study are available at https://github.com/RJauss/ParasitesParadise.

## Author contributions

MB and MS conceived the study. RW and StS designed the sampling and DNA extraction. AMF-D contributed the primers and functional annotation of oomycetes. SW and R-TJ conducted the sampling, DNA extraction and PCRs. KF assisted DNA extraction and PCRs. RT-J performed the bioinformatic and statistical analyses and drafted the manuscript. All authors contributed to and approved the final version.

## Supplementary Figures

**Supplementary Figure 1: Sequence similarity of reads (top) and OTUs (bottom) per sampling event to published reference sequences.** 20.5% of all OTUs, corresponding to 3% of all reads, had a similarity of less than 70% to any known reference sequence (not shown).

**Supplementary Figure 2: Taxonomic assignment of OTUs per sampling and microhabitat.** Black line separates canopy and ground habitats. Distribution of taxonomic groups was similar for every sampling, i.e. Pythiales and Peronosporales dominating all samples.

**Supplementary Figure 3: Ternary plot partitioning the relative abundances of Peronosporales and Pythiales per sampling event.** Each dot represents one OTU.

**Supplementary Figure 4: Differential abundance analysis between the two seasons spring (top panels) and autumn (bottom panels) sorted by taxonomic order.** Each dot represents one significantly differentially abundant OTU grouped by genus. Y-axis (log2FoldChange) gives the measurement of the differential abundance.

**Supplementary Figure 5: Boxplot of alpha diversity indices for microhabitat communities per sampling.** Outliers are given by dots. Observed patterns show no strong variability over the four sampling events.

## Supplementary Tables

**Supplementary Table 1: Primer tags used in this study.** Given are the sample ID, forward and (reverse complemented) reverse tag and the ENA sequencing run ID.

